# Cooperative stator assembly of bacterial flagellar motor mediated by rotation

**DOI:** 10.1101/2020.04.26.059089

**Authors:** Kenta I. Ito, Shuichi Nakamura, Shoichi Toyabe

## Abstract

Cooperativity has a central place in biological regulation, providing robust and highly-sensitive regulation. The bacterial flagellar motor (BFM) implements autonomous torque regulation based on the stator’s nonequilibrium structure; the stator units bind to and dissociate from the motor dynamically in response to environmental changes. However, the mechanism of this dynamic assembly is not fully understood. Here, we demonstrate the cooperativity in the stator assembly dynamics. The binding is slow at the stalled state, but the rotation in either direction boosts the stator binding. Hence, once a stator unit binds, it drives the rotor and triggers the avalanche of succeeding bindings. This cooperative mechanism based on nonequilibrium allostery accords with the recently-proposed gear-type coupling between the rotor and stator.

---

Motility regulation is vital and the basis of autonomy for many life forms. Swimming bacteria like *Escherichia coli* and *Salmonella enterica* have been serving as the model system of the motility regulation. Since they utilize the propulsive force of the rotating flagella, the torque regulation of the flagellar motor, as well as the control of tumbling frequency and flagellar bundle formation, is essential for the motility regulation. The bacterial flagellar motor (BFM)^1–4^ (Fig. 1A) is a large protein complex and can rotate at up to 1,700 Hz^5^, reverse the rotation, and vary the torque depending on the load on the motor. A striking fact is that the BFM has multiple torque-generating stator units and regulates the torque magnitude by dynamically alternating the units between the motor and surrounding membrane pool^6–8^ (Fig. 1B). This autonomous stator assembly relies on the self-regulation of each stator unit. The stator unit implements a load sensor and a load-dependent regulator^9–12^; the stator-unit dissociation is suppressed at high motor load for keeping large torque generation and enhanced at low load for probably reducing futile energy consumption. On the other hand, the binding rate is thought to be not significantly dependent on the load^11^. The ion flux through the ion channel of the stator unit is required for the stator-unit binding^13–15^. However, it is not clear how the ion flux is involved in the binding process.

**Figure 1.**
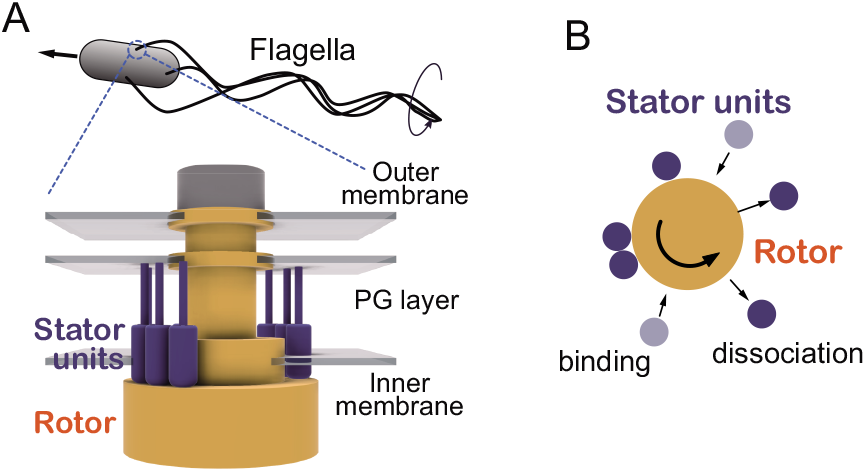
**A**, The bacterial flagellar motor is a supramolecular complex with a diameter of about 45 nm spanning the membrane^1–4^. The motor consists of a rotor (yellow) and multiple torque-generating stator units (blue). The stator units have ion channels and generate torque at the interface with the rotor by exploiting the free energy change liberated by the ion currents through the channels. The stator units interact with the rotor at its bottom part and are anchored to the rigid peptidoglycan (PG) layer with its top part^16–18^. **B**, The stator units bind to and dissociate from the motor.

Cooperativity is central to the biological regulation^19^. A simple allosteric mechanism provided by the mutual dependence of the ligand-binding sites exhibits a steep nonlinear response, enabling robust and highly-sensitive regulation and signaling^20^. It would be natural to suppose that the BFM’s torque regulation is highly optimized. In this work, we demonstrate the cooperative stator assembly mediated by rotation. The stator binding is slow at the stalled state, but the rotation in either direction boosts the stator binding. Since the binding of the first stator unit causes the rotation, there is a cooperative feedback loop of the stator assembly via rotation.

We exploit the dynamic load control by the electrorotation to study the stator remodeling process^12,21^. The electrorotation induces a rotating high-frequency (~MHz) electric field to apply external torque on microscopic dielectric objects such as the bacterial cell body. Hence, we can induce torque on the motor by tethering the flagellum to a glass surface. Since the torque magnitude is proportional to the electric field’s squared amplitude 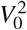, we can modulate the torque dynamically.

## Results

### Stator remodeling

Each experimental cycle consists of initial, assisting, and remodeling phases (Fig. 2A). In the initial phase, we observed a steady rotation for 10 s in the absence of an external load. The number of the bound stator units, *N*, was estimated to be approximately ten in this phase by the step analysis in below. This relatively large value of *N* is due to the high viscous load on the tethered cell and is consistent with the previous results^21^.

**Figure 2.**
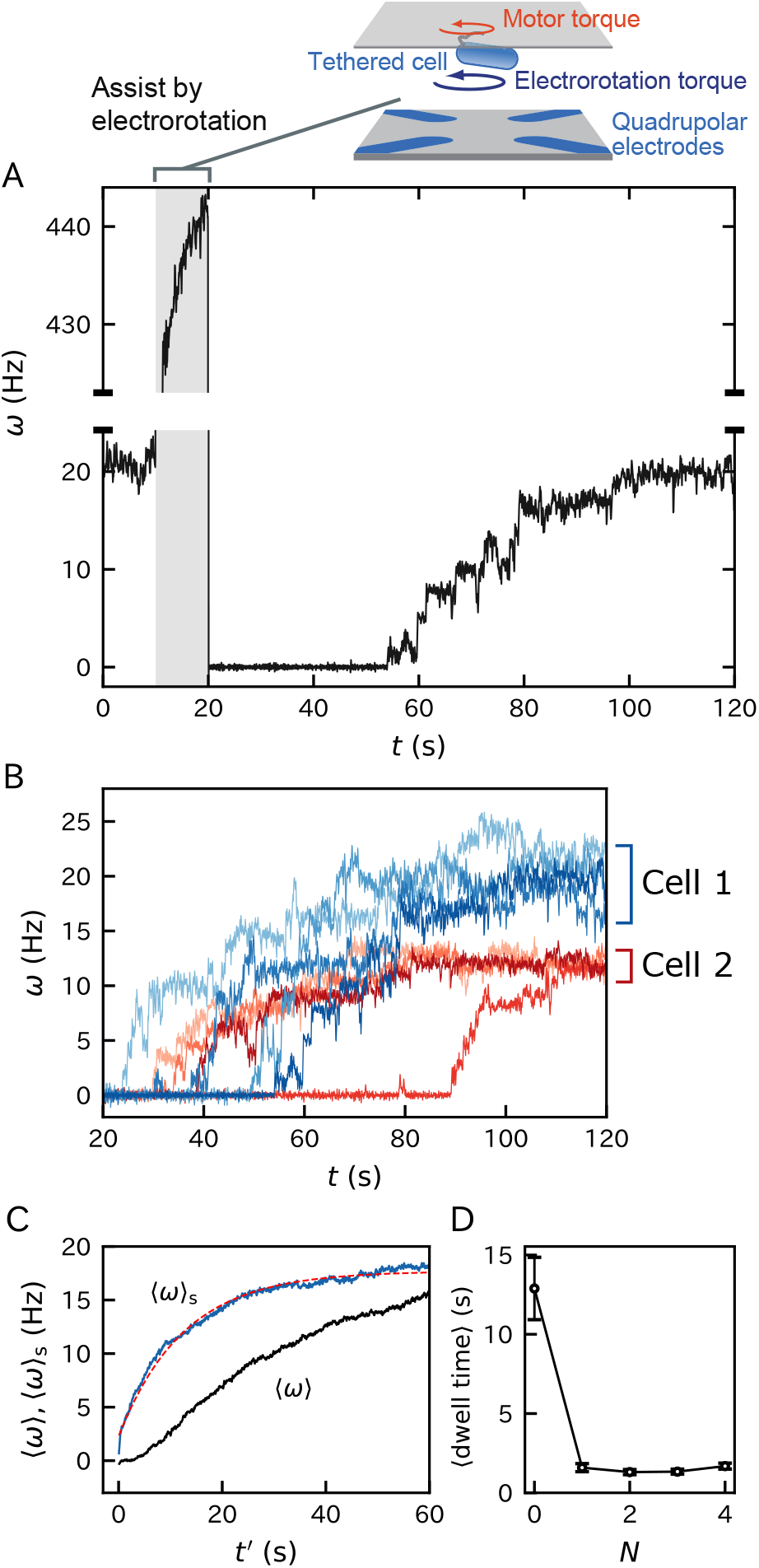
Rotation traces. **A**, A typical rotation trace of the remodeling process. After the initial 10-s phase, a strong assisting torque was applied for 10 s for releasing the stator units from the motor by the electrorotation method^21^. *ω* > 0 corresponds to the counter-clockwise (CCW) direction. **B**, Consecutive traces of two motors are superposed. **C**, Ensemble average of remodeling traces including *N* = 0 state (〈*ω*〉, black) and that excluding it (〈*ω*〉_s_, blue). 40 traces were averaged. For 〈*ω*〉, the time origin was reset to *t*′ = 0. 〈*ω*〉_s_ was fitted by 〈*ω*〉_s_ = *ω*_ss_(1 − *e^−kt′^*) + *ω*_1_*e^−kt′^* with the fitting parameters of *k* = 0.080s^−1^ and *ω*_ss_ = 18s^−1^(red dashed curve). *ω*_1_ was approximated by *ω*_max_/10. **D**, The mean dwell time at each *N* obtained by the step analysis of the traces (Fig. S1). The error bars indicate standard errors.

In the assisting phase, we assisted the rotation strongly for 10 s by the electrorotation for reducing the load on the motor. The rotation speed was accelerated to about 300 - 600 Hz depending on the body dimension and cellular dielectric property.

In the remodeling phase, the assisting torque was turned off. The rotation rate vanished in the most traces (about 80% of the observed traces). The motor stalling indicates that all the bound stator units are released under the low-load condition^12, 21^. Some motors continued rotating while their rotation rate decreased much compared to the initial phase. This may be due to the insufficient magnitude and duration of the assisting torque.

The rotation rate recovered to the initial steady level with successive jumps, indicating the stepwise stator-unit bindings. The duration of each cycle was 120 s. We repeated multiple cycles for the same motor.

### Slow binding of first stator unit

We found strong stochasticity in the traces of the rotation rate *ω*, especially at small *ω* (Fig. 2B). The duration of the zero-speed state was typically long compared to the duration of the other states. That is, the binding rate of the first stator unit is small. However, once a stator unit binds, it triggers the bindings of succeeding stator units.

The slow binding at *N* = 0 is highlighted when we compared the ensemble average of the traces including *N* = 0 state (〈*ω*〉) and that excluding it (〈*ω*〉_s_) (Fig. 2C). For 〈*ω*〉_s_, we excluded *N* = 0 state by shifting each trace so that the first stator unit bound at *t*′ = 0. We located the binding as *ω* exceeded *ω*_c_ ≡ *ω*_max_/20. *ω*_max_ is the maximum rotation rate except for the duration where external torque is induced. Since approximately ten stator units bind at the steady state, *ω*_c_ corresponds to the half of the rotation rate caused by a single stator unit. When the traces intersected *ω*_c_ multiple times, we used the last one. We found that 〈*ω*〉_s_ deviated significantly from 〈*ω*〉. 〈*ω*〉 showed a downward convexity at small *t*′ in contrast to the upward convexity of 〈*ω*〉_s_. This characteristic is expected to be caused by the long duration of *N* = 0 state.

A theoretical curve, 〈*ω*〉_s_ = *ω*_ss_(1 − *e^−kt′^*) + *ω*_1_*e^−kt′^*, fitted 〈*ω*〉_s_ well with the fitting parameters of the steady-state rotation rate *ω*_ss_= 18 s^−1^ and the rate constant *k* = 0.080s^−1^ (Fig. 2C). *ω*_1_ is the rotation rate caused by a single stator unit and was approximated by *ω*_max_/10. This curve is the solution of the random packing model^11^ given by *d(ω_s_/dt’ = k*(*ω*_ss_ - (*ω*_s_) under the initial condition of (*ω*_s_(0) = *ω*_1_. *k* is the sum of the binding rate at a binding site *k*_+_ and the dissociation rate of a bound stator *k*_-_. The successful fitting may validate the model assumption for *N* ≥ * 1; *k*_±_ are independent of *N*.

The step analysis of the remodeling traces (Fig. S1) supported the long pausing at *N* = 0 compared to those at *N* ≥ 1 (Fig. 2D). This implies that *N* = 0 is a metastable state.

Additional experiments were done to eliminate the possibility that the slow binding at *N* = 0 is caused by an experimental artifact. The possible concerns include the cell-surface interaction and the cellular damage caused by the temperature rise by the electrorotation. As the test, we observed remodeling traces under degraded condition. We used cells without a shearing treatment, blocking agent with reduced concentration (50 mg/ml), and observation buffer containing high ion strength (10 mM MOPS and 10 mM KCl). With these modifications, the cells can easily stick to the chamber surface, and the temperature increased three times as much as the present condition during the assisting phase (Fig. S2). Nonetheless, we obtained quantitatively similar remodeling traces and dwell time (Fig. S3). This verifies that the slow binding at *N* = 0 is the motor characteristic.

### Rotation enhances binding

For elucidating the mechanism of the slow binding at *N* = 0, we investigated the remodeling process under forced rotation (Fig. 3). We calibrated the rotation rate by dividing by the steady-state rotation rate during the intitial phase *ω*_ss_. The assisting phase was followed by a 1-s duration without external torque for verifying that all the stator units are released. We analyzed only the trajectories with vanished rotation rate. Then, a constant external torque was applied for 15 s. Accordingly, steep change in *ω* was observed at the start and end of the torque induction. These jumps had similar magnitudes and opposite signs as expected.

**Figure 3.**
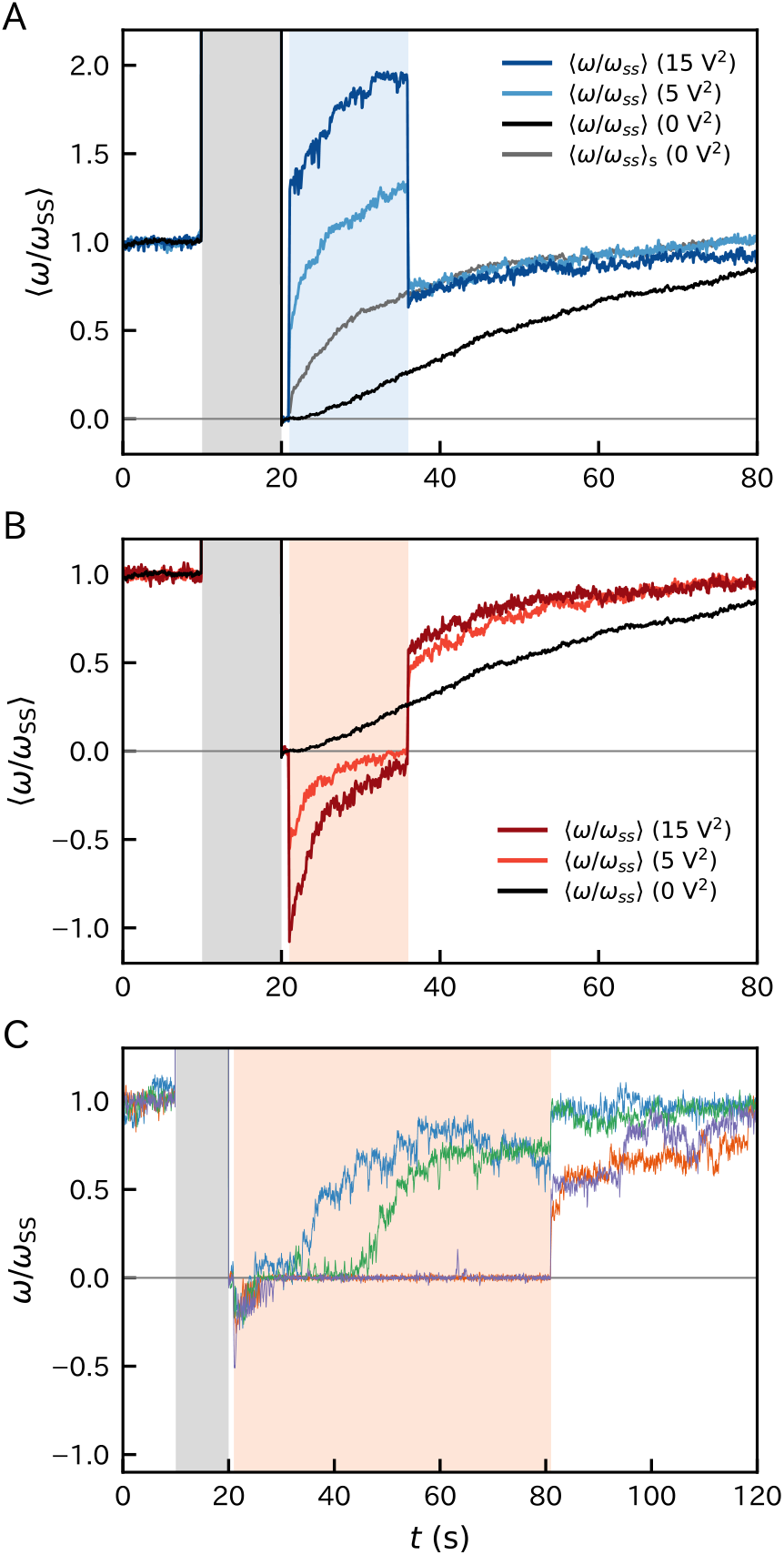
Ensemble averages of the remodeling traces under constant external torque in the assisting direction (**A**) and hindering direction (**B**) with different 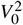. The torque was induced for 15 s (**A, B**) or 60 s (**C**) starting at 21 s. The rotation rate *ω* was first normalized by the steady rotation rate *ω*_ss_ calculated as the mean *ω* during the initial phase and then averaged. The black and gray curves are the calibrated curves of 〈*ω*〉 and 〈*ω*〉_s_ of Fig. 2C. For (*ω*)_s_, the time origin was shifted to *t* = 21 s. **C**, The remodeling process with extended torque induction in the hindering direction.

Under the forced rotation in either direction, the remodeling proceeded quickly with upward convexity (Fig. 3A, B). As well, at the timing when assisting torque was turned off, the values of 〈*ω/ω*_ss_〉 under forced rotation were larger than that without forced rotation (Fig. 3A). That is, more stator units bound with the forced rotation, suggesting that the rotation enhances the binding. The curves under different external torque as well as 〈*ω*/*ω*_ss_〉_s_ were similar except for the bias during the torque induction and coincided well after the torque was turned off. This indicates that the binding dynamics does not significantly depend on the magnitude of the external driving.

Interestingly, the remodeling dynamics under the assisting and hindering torque were asymmetric. When assisted, the remodeling continued until *ω* recovers to the level of the initial phase (Figs. 3A and S4). On the other hand, when hindered, the remodeling started quickly as well but stalled when *ω* reached zero (Figs. 3B, C, and S4). The stall typically continued until the removal of the induced torque, and then the remodeling restarted quickly. This characteristic did not change qualitatively with a different magnitude of hindering torque. Further remodeling during the stall was occasionally observed when we extended the duration of torque induction (Fig. 3C). These results suggest that the binding is slow even for *N* ≠ 0 if |*ω*| vanishes and is enhanced by the rotation in either direction.

During the stall under hindering, the motor torque *T*_m_ and electrorotation torque *T*_ex_ should be balanced. However, this is not expected considering the discrete nature of *T*_m_. It is expected that *T*_m_ takes discrete values proportional to *N* under the present high load condition of the tethered cell assay. That is, the perfect balance between *T*_m_ and *T*_ex_ is not expected. Nevertheless, the remodeling under hindering torque experienced the stall in most trajectories. This is probably due to the pinning of the cell caused by, for example, tiny obstacles on the glass surface.

We observed that, when the cells are forcedly rotated at a slow rate of ~ 1 Hz, the further binding did not proceed quickly (Fig. S4A). The binding enhancement by rotation may have a threshold for *ω*. However, we had difficulty in determining the threshold value since the slow rotations suffered from the cell-surface interaction and stopped occasionally.

### Quantitative evaluation of binding enhancement under forced rotation

We evaluated the binding enhancement under forced rotation. We exploited pulse-wise external torque instead of the constant torque (Fig. 4A, B). This is because the binding may occur very soon after the torque induction. If this is the case, it is difficult to distinguish the increase in *ω* by the torque induction and that by the stator-unit binding.

**Figure 4.**
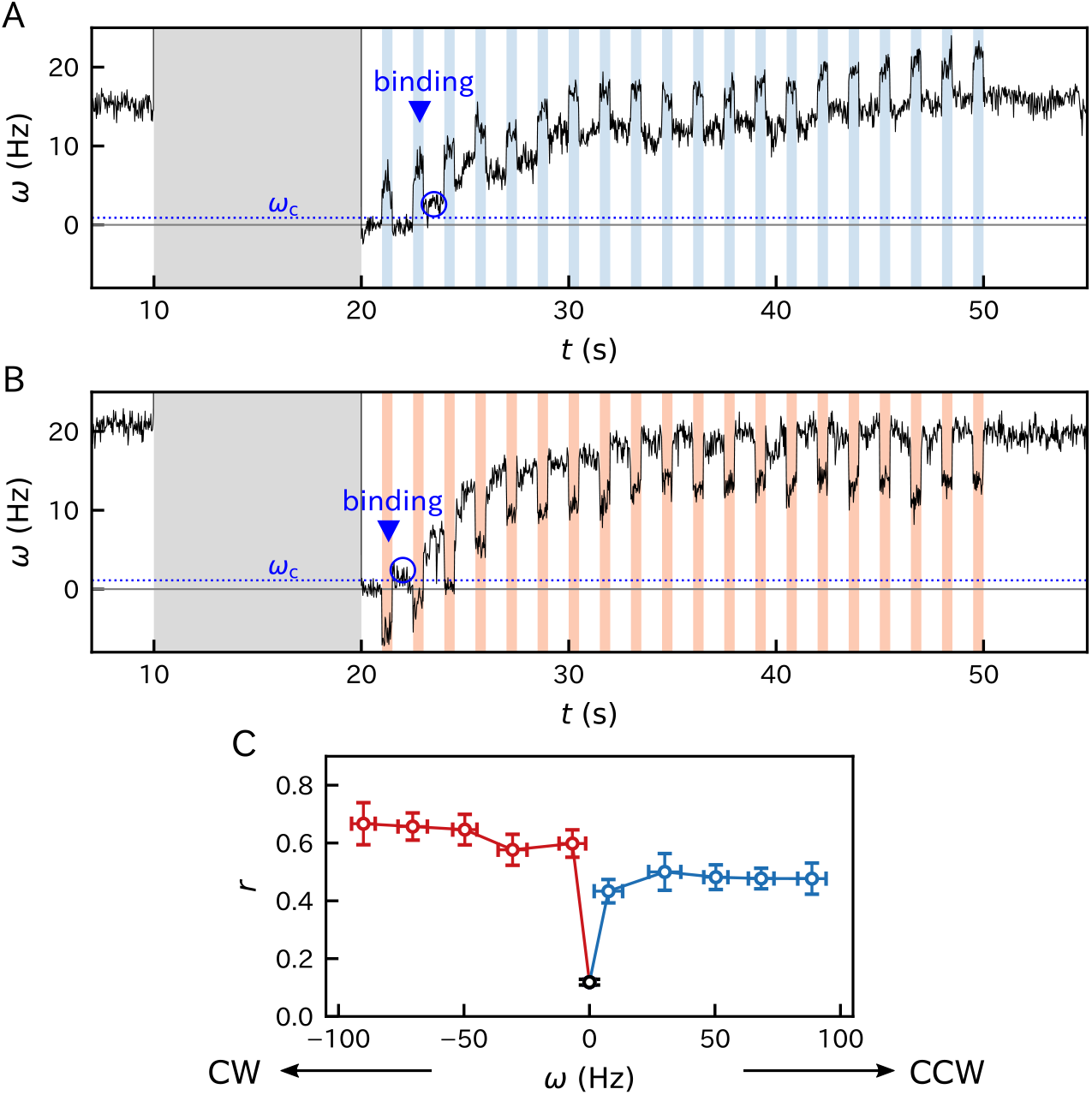
Evaluattion of rotation-dependent binding rate using pulse-wise forced rotation. The alternation of 0.5-s ON (shaded by blue or red) and 1-s OFF of the external torque was induced in assisting direction (**A**) or hindering direction (**B**). When *ω* exceeded *ω*_c_ ≡ *ω*_max_/20, we assumed that the first stator unit bound. **C**, Binding rate as a function of the forced rotation rate averaged in bins with the width of 20 Hz. We used 396 traces of 122 cells in total. See Fig. S5 for details. The error bars indicate the standard errors.

We repeated short torque induction (ON) lasting for *τ* = 0.5 s with a one-second break (OFF) during the remodeling phase. We assumed that the first stator unit bound during an ON when the mean of *ω* in the succeeding OFF exceeded *ω*_c_. We counted the number of binding events (*N* = 0 → 1), *n*_+_, and non-binding events (*N* = 0 → 0), *n*_0_, at *N* = 0. These numbers were accumulated for multiple traces. We used 396 traces of 122 cells in total. The fraction *r* = *n*_+_/(*n*_+_ + *n*_0_) quantifies how the forced rotation enhances the binding.

The forced rotation in either direction significantly enhanced the binding (Fig. 4C). *r* depended on the rotational direction and was larger for the clockwise (CW) rotation than for the counter-clockwise (CCW) rotation. A significant dependence of *r* on *ω* was not observed. Specifically, we could not resolve the gradual increase in *r* at small *ω*. The plots with varied bin widths indicated that the threshold value of |*ω*| for the binding enhancement was estimatedto be less than ~ 3 Hz (Fig. S5).

Note that the binding rate at a single binding site is approximated by 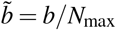 with *b* = −*τ*^−1^ ln(1 − *r*). Here, the bound stator unit is supposed not to dissociate during the same ON. The probability that an event obeying a Poissonian process with the rate *b* takes place within *τ* is *r* = 1 − exp(−*bτ*), which leads to the above relation. The average of *b* in the CCW rotation was 1.3 ± 0.1s^−1^ (Fig. S5), which yields 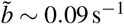 assuming *N*_max_ = 14. On the other hand, the fitting in Fig. 2C provided *k = k_+_ + k*_−_ = 0.080s^−1^, which corresponds to the rate constant under the CCW rotation driven internally by the stator units. For the tethered cells, *k ≃ k*_+_ is expected since *k*_−_ is negligible at high load^11^. Hence, the similarity of *k* and 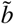 implies that, at least in the CCW rotation, the binding rate is solely determined by the rotation rate independent of whether the rotor is externally rotated or internally rotated by the stator units.

## Discussion

### Cooperative stator assembly via rotation

We evaluated the stator-assembly dynamics during the remodeling process based on the dynamical load control provided by the electrorotation. The stalled state (*ω* = 0) is metastable, and the binding of the stator unit to the stalled rotor is slow. However, the binding is significantly enhanced by the rotation in either direction with |*ω*| exceeding a small threshold. The CW rotation was more effective than the CCW rotation. The binding of a stator unit induces the rotation and, therefore, triggers the succeeding stator-unit bindings like an avalanche. Thus, the nonequilibrium allostery via rotation mediates the cooperativity of the stator assembly. The evaluation of the threshold values and the binding and dissociation rate under forced rotation remains for future studies.

The ion flux is required for the stator assembly^13–15^. Without the ion flux, the stator units do not generate torque and rotate the rotor. This is consistent with our result.

It was previously shown that the binding rate does not depend on the load at intermediate *N*^11^, while the dissociation rate has a significant dependence on the load^11,12^. We showed that the binding is enhanced in either rotational direction despite that the load on the stator is different between them; small or negative with the assisting torque and large with the hindering torque. This supports that the binding is regulated not by the load even at small *N* but by the rotation rate. It is natural that the stator unit that is not yet incorporated does not feel the load. A question is how the stator unit can sense the rotation.

### Activation by rotation

The structure of the stator unit implies that the stator unit itself is a rotating motor. The MotA5 ring may revolve around the MotB2 axis and rotate the rotor via a gear-type coupling^22–24^ (Fig. 5). Such coupling may explain our finding that the rotor rotation accelerates the binding of the stator unit as follows.

**Figure 5.**
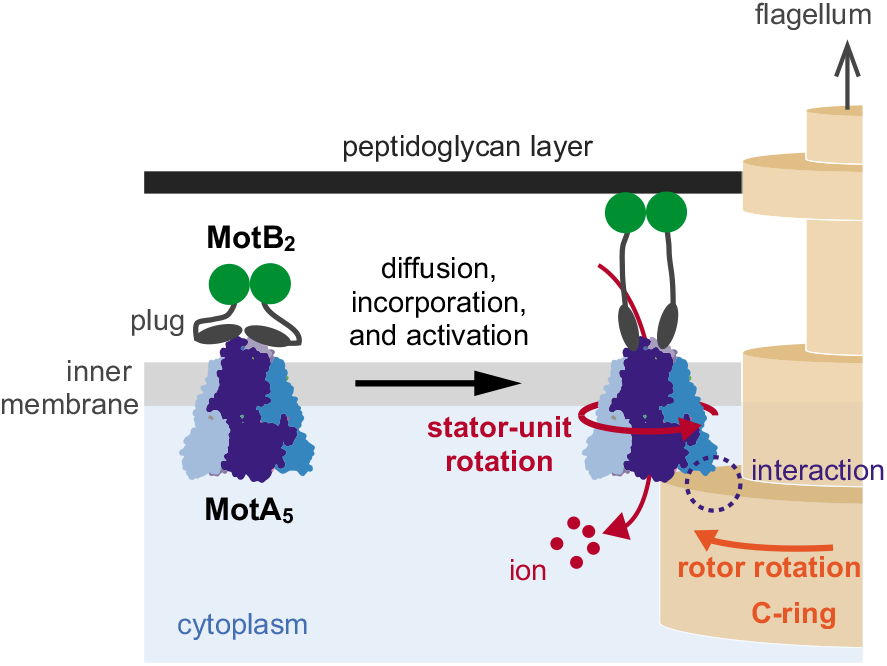
The binding process of the stator unit to the rotor. The freely-diffusion stator unit collides with the rotor. The electrostatic interaction with the C-ring induces a conformational change of the stator unit, which allows the ion flow and, hence, the torque generation.

The stator-unit binding would involve the interaction between the stator unit and rotor^25^and the binding of MotB’s C-terminal to the peptidoglycan layer as an achor^16–18^. This mechanism remains elusive. A hypothesis is that the stator unit diffusing in the membrane pool collides with the motor, binds partially to the rotor, and will be activated. Then, the stator unit starts torque generation. Although this activation process is not known clearly, the gear-type coupling may give a hint. The rotor rotation may rotate the stator unit via the gear and induce the stator unit’s conformational change necessary for the activation. It is natural to assume that such the rotation-mediated activation can be differently induced by the CW and CCW rotation (Fig. 4C).

### Function of rotation-mediated cooperativity

The metastability of the stalled state may be beneficial for bacteria by ensuring that the rotor does not rotate during the motor assembly and that futile bindings more than necessary do not occur when stacked in biofilm. More studies would clarify the function served by the binding enhancement by rotation.

An intriguing question related to the function is whether a rotational fluctuation enhances the binding or unidirectional rotation is necessary. The motor under the swimming condition is expected to have larger rotational fluctuations at a stall than those of the tethered cell. The determination of the threshold rotation rate for the binding enhancement may provide a hint for answering this question.

## Acknowledgements

We thank Yoshiyuki Sowa, Yohei Nakayama, and Kazuo Sasaki for helpful discussions. This work was supported by JSPS KAKENHI (16H00791 and 18H05427).

## Methods

### Cell preparation

*Salmonella* strain YSC2123, which lacks *motA, motB, cheY, fimA*, and *fliC* (204–292), was transformed with a plasmid encoding wild-type *motA/motB*^26^ (referred to as a wild-type). This strain has a “sticky” filament with a hydrophobic surface beneficial for the tethered cell assay. Cells were grown in L-broth containing 100 μg/ml ampicillin and 0.2% arabinose (for the expression of the MotA and MotB) for at least 10 hours at 30 °C with shaking. The buffer was replaced with the observation buffer (5mM MOPS(3-Morpholinopropanesulfonic acid) and 5 mM KCl adjusted to pH8.3 with KOH). We partially sheared the flagella filaments by passing the bacterial solution through 25G needle 70 times.

### Microscopy

We observed the rotation of a tethered cell at room temperature (24 °C) on a phase-contrast upright microscope (Olympus BX51WI, Japan) with a 60× objective lens (Olympus, NA=1.42), at 4,000 Hz using a high-speed CMOS camera (Basler, Germany), high-intensity LED (623 nm, 4.8W, Thorlabs, NJ) for illumination, and a laboratory-made capturing software developed on LabVIEW (National Instruments, TX). The angular position of the cellular body was analyzed by an algorithm based on a principal component analysis of the cell image. We used 250mg/ml Perfect Block (MobiTec, Germany) as the blocking agent to suppress the interaction between the cell body and the glass surface. The chamber height was about 20 μm.

### Electrorotation

A 10-MHz sinusoidal voltage with a phase shift of *π*/2 was induced on the four electrodes patterned on the bottom glass surface (Fig. 1A). The distance between the electrodes is 47 μm. The signal was generated by a function generator (nf, Japan) controlled by PC and divided by 180° phase distributors (Thamway, Japan). They were amplified by four amplifiers (Analog Devices, MA) and loaded on the electrodes. This generates an electric field rotating at 10 MHz in the center of the electrodes and induces a dipole moment rotating at 10 MHz on the cell body. Since there is a phase delay between the electric field and the dipole moment, the cell body is subjected to a constant torque. The torque magnitude is proportional to the square of the voltages’ amplitude *V*_0_. We modulated *V*_0_ by a signal generated by the multifunction board (National Instruments) equipped on PC. The camera and amplitude signal were synchronized at a time difference of less than one microsecond. We applied an external torque with a time-dependent magnitude, as indicated in the main text, with superposed by a 1000-Hz small sinusoidal torque for the possible torque calibrations^21, 27^ although we did not calibrate torque in this work.

The temperature under electrorotation was measured using a pair of temperature-dependent fluorescent dyes, Rhodamine B and Rhodamine 101, with different temperature dependence^28^. The ratio of the fluorescence of these dyes provide the temperature. The estimated temperature during electrorotation nwas 28.1 °C for 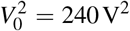, which is the typical value used during the assisting phase, at the room temperature of 24 °C (Fig.S2). The temperature relaxed to room temperature soon within a second after the electrorotation was turned off.

### Step analysis

We estimated the dwell time at small *N* during the remodeling process by a step analysis algorithm developed recently^10,12^.(Fig. 2D). The trajectories are typically noisy. We analyzed 40 remodeling traces (19 cells) with seemingly clear steps. Approximately the same number of erroneous traces was not used.

The algorithm tries to find the large velocity jump in the traces by comparing the mean velocities of adjacent segments of the traces. The core part of the algorithm is briefly summarized in the following. The traces were first averaged with moving windows with the length of 500 frames and 100-frame shift. Then, the traces were divided into small segments with the length of 5 points. The adjacent segments were merged when the mean velocities were smaller than a threshold value, *ω*_max_/15. Here, *ω*_max_ is the maximum of the moving-averaged *ω* during the initial phase and also the last part of the remodeling phase (*t* ≥ 50s). This procedure was iterated until adjacent segments could no longer be merged. Then, the segment boundaries were adjusted so that the *χ*^2^ for the two segments are minimized. If this adjustment generates a segment shorter than 5 points, the segment was merged to the adjacent segment.

### Binding rate

Let *t_i_* be the starting time of the *i*-th OFF state. We assumed that the stator unit bound in the *i*-th ON state if 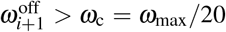. Here, 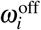 is the mean rotation rate during *t_i_* + 0.05s ≤ *t < t_i_* + 0.15s. This limitation avoids to include the relaxation right after the torque switching and also the possible binding during the OFF state. Since approximately 10 stator units are thought to be bound at maximum under the present condition, *ω*_c_ corresponds to the half of the rotation rate caused by a single stator unit.

When the binding takes places in the *i*-th ON state, the corresponding rotation rate is determined as the mean rotation rate during *s_i_* + 0.05s ≤ *t ≤ s_i_* + 0.15s. Here, *s_i_* is the starting time of the *i*-th ON state.

## Data availability

The data that support the findings of this study are available from the corresponding author upon reasonable request.

## Code availability

The computer codes used for this study are available from the corresponding author upon reasonable request.

## Author Contributions

KI, SN and ST designed research and wrote the paper. KI performed experiments. KI and ST contributed analytic tools and analyzed data.

## References

1. Berg, H. C. The rotary motor of bacterial flagella. Annu. Rev Biochem. 72, 19–54 (2003).

2. Berry, R. M. & Armitage, J. P. The bacterial flagella motor. Adv Microb Physiol. 41, 291–337 (1999).

3. Sowa, Y. & Berry, R. M. Bacterial flagellar motor. Q. Rev. Biophys. 41, 103–132 (2008).

4. Nakamura, S. & Minamino, T. Flagella-driven motility of bacteria. Biomolecules 9, 279 (2019).

5. Magariyama, Y. et al. Very fast flagellar rotation. Nature 371, 752–752 (1994).

6. Block, S. M. & Berg, H. C. Successive incorporation of force-generating units in the bacterial rotary motor. Nature 309, 470–2 (1984).

7. Ryu, W. S., Berry, R. M. & Berg, H. C. Torque-generating units of the flagellar motor of escherichia coli have a high duty ratio. Nature 403, 444 (2000).

8. Leake, M. C. et al. Stoichiometry and turnover in single, functioning membrane protein complexes. Nature 443, 355 (2006).

9. Tipping, M. J., Delalez, N. J., Lim, R., Berry, R. M. & Armitage, J. P. Load-dependent assembly of the bacterial flagellar motor. mBio 4, 00551–13 (2013).

10. Lele, P. P., Hosu, B. G. & Berg, H. C. Dynamics of mechanosensing in the bacterial flagellar motor. PNAS 110, 11839–44 (2013).

11. Nord, A. L. et al. Catch bond drives stator mechanosensitivity in the bacterial flagellar motor. PNAS 114, 12952–12957 (2017).

12. Wadhwa, N., Phillips, R. & Berg, H. C. Torque-dependent remodeling of the bacterial flagellar motor. PNAS 201904577 (2019).

13. Fukuoka, H., Wada, T., Kojima, S., Ishijima, A. & Homma, M. Sodium-dependent dynamic assembly of membrane complexes in sodium-driven flagellar motors. Mol. Microbiol. 71, 825–835 (2009).

14. Tipping, M. J., Steel, B. C., Delalez, N. J., Berry, R. M. & Armitage, J. P. Quantification of flagellar motor stator dynamics through in vivo proton-motive force control. Mol. Microbiol. 87, 338–347 (2013).

15. Sowa, Y., Homma, M., Ishijima, A. & Berry, R. M. Hybrid-fuel bacterial flagellar motors inEscherichia coli. PNAS 111, 3436–3441 (2014).

16. Kojima, S. et al. Stator assembly and activation mechanism of the flagellar motor by the periplasmic region of MotB. Mol. Microbiol. 73, 710–718 (2009).

17. Terahara, N. et al. Na+-induced structural transition of MotPS for stator assembly of the Bacillus flagellar motor. Sci. Adv. 3, eaao4119 (2017).

18. Kojima, S. et al. The helix rearrangement in the periplasmic domain of the flagellar stator b subunit activates peptidoglycan binding and ion influx. Structure 26, 590–598.e5 (2018).

19. Williamson, J. R. Cooperativity in macromolecular assembly. Nat. Chem. Biol. 4, 458–465 (2008).

20. Pauling, L. The oxygen equilibrium of hemoglobin and its structural interpretation. PNAS 21, 186–191.

21. Sato, K., Nakamura, S., Kudo, S. & Toyabe, S. Evaluation of the duty ratio of the bacterial flagellar motor by dynamic load control. Biophys. J. 116, 1952–1959 (2019).

22. Santiveri, M. et al. Structure and function of stator units of the bacterial flagellar motor. Cell 183, 244–257.e16 (2020).

23. Deme, J. C. et al. Structures of the stator complex that drives rotation of the bacterial flagellum. Nat. Microbiol. (2020).

24. Chang, Y. et al. Molecular mechanism for rotational switching of the bacterial flagellar motor. Nat. Str. Mol. Biol. 27, 1041–1047 (2020).

25. Morimoto, Y. V., Nakamura, S., Kami-ike, N., Namba, K. & Minamino, T. Charged residues in the cytoplasmic loop of MotA are required for stator assembly into the bacterial flagellar motor. Mol. Microbiol. 78, 1117–1129 (2010).

26. Morimoto, Y. V., Che, Y. S., Minamino, T. & Namba, K. Proton-conductivity asssay of plugged and unplugged MotA/B proton channel by cytoplasminc pHluorin expressed in Salmonella. FEBS Lett. 584, 1268–72 (2010).

27. Toyabe, S., Watanabe-Nakayama, T., Okamoto, T., Kudo, S. & Muneyuki, E. Thermodynamic efficiency and mechanochemical coupling of F1-ATPase. PNAS 108, 17951–17956 (2011).

28. Sakamoto, Y. & Toyabe, S. Assembly of a functional and responsive microstructure by heat bonding of DNA-grafted colloidal brick. Sci. Rep. 7 (2017).

