## Supplemental Information for "Cooperative stator assembly of bacterial flagellar motor mediated by rotation"

### Supporting Information for "Cooperative stator assembly of bacterial flagellar motor mediated by rotation"

#### S1. Step analysis

Figure S1 plots all the traces and the step-analysis used for Fig. 2.

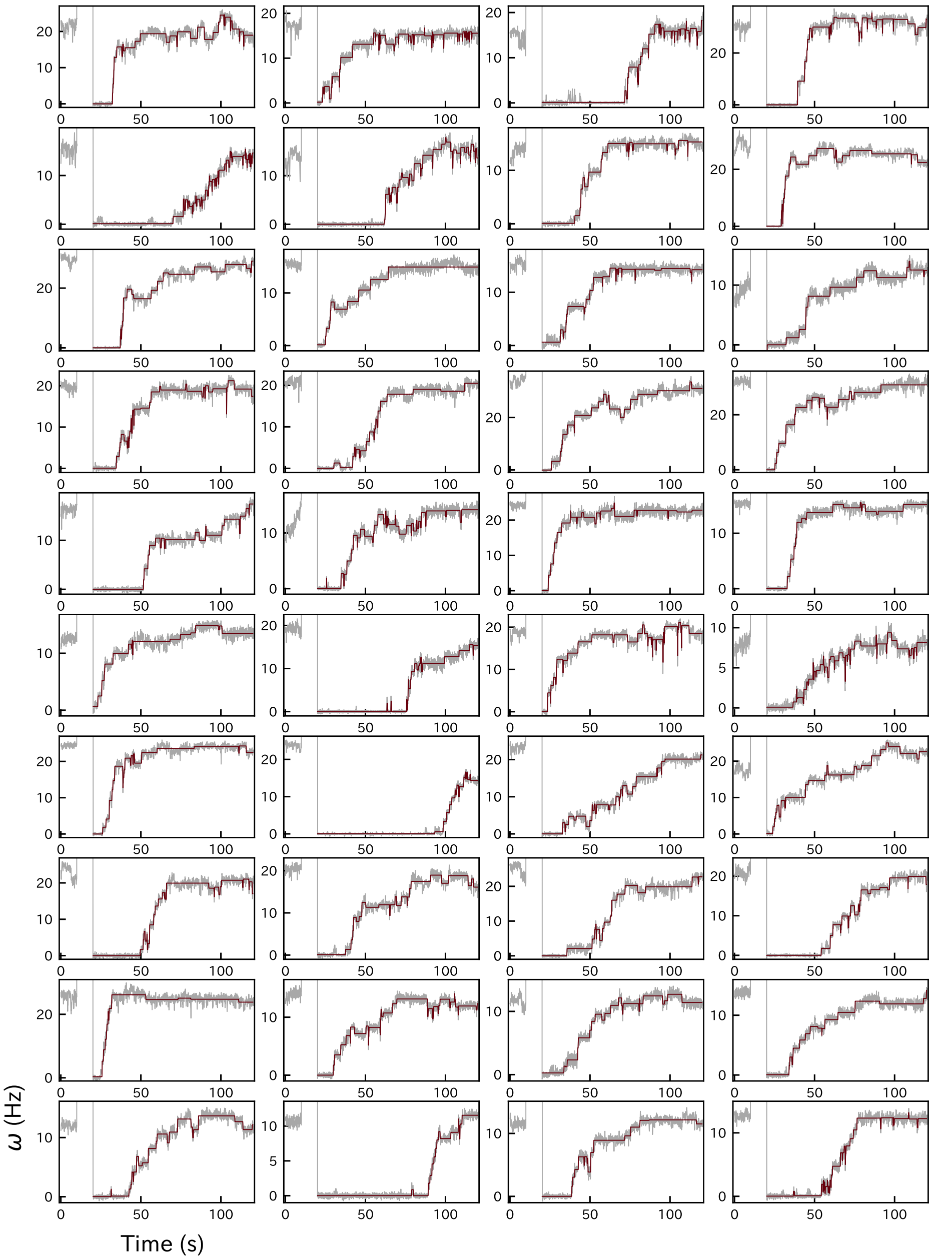


**Figure S1.** The traces used for Fig. 2. The rotation traces smoothed by the moving average with the window length of 500 points and frame shift of 100 points (gray) are superposed by the step-analysis traces (red).

#### S2. Temperature rise

The temperature under electrorotation was measured using two fluorescent dyes, Rhodamine B (Sigma-Aldrich) and Rhodamine 101 (Sigma-Aldrich). These dyes have fluorescence with different dependence on the temperature. The spatial temperature gradient may induce a thermophoretic migration of the dyes. However, the ratio of the two dyes provides the temperature assuming that the thermophoretic magnitude is the same for these two dyes. The calibration curve was measured using a realtime thermal cycler (BioRad). The electrorotation chamber was first filled with 50 μg/ml Rhodamine B, 50 μg/ml Rhodamine 101, or observation buffer without fluorescent dye were flowed into the chamber. The fluorescence of these three solutions with/without electrorotation were measured at 16 Hz and analyzed to estimate the temperature profile (Fig. S2).

The temperature rise by electrorotation in the observation buffer containing 5 mM MOPS and 5 mM KCl, which was the standard one int this work, was estimated to be 1.8 °C for $V_0^2 = 100\,\mathrm{V}^2$, 3.5 °C for 200 $\mathrm{V}^2$, and 5.1 °C for 300 $\mathrm{V}^2$ (Fig. S2A, B, C). The temperature relaxed to the room temperature soon within a few seconds after the electrorotation was turned off. The typical value of $V_0^2$ during the assisting phase was 240 $\mathrm{V}^2$, which yields the temperature increase of 4 °C by interpolating the above values.

On the other hand, in a high-ion-strength buffer containing 10 mM MOPS and 10 mM KCl used for the test (Fig. S3), the temperature raised approximately 12 °C for 300 $\mathrm{V}^2$ (Fig. S2F), which was the typical voltage in Fig. S3.

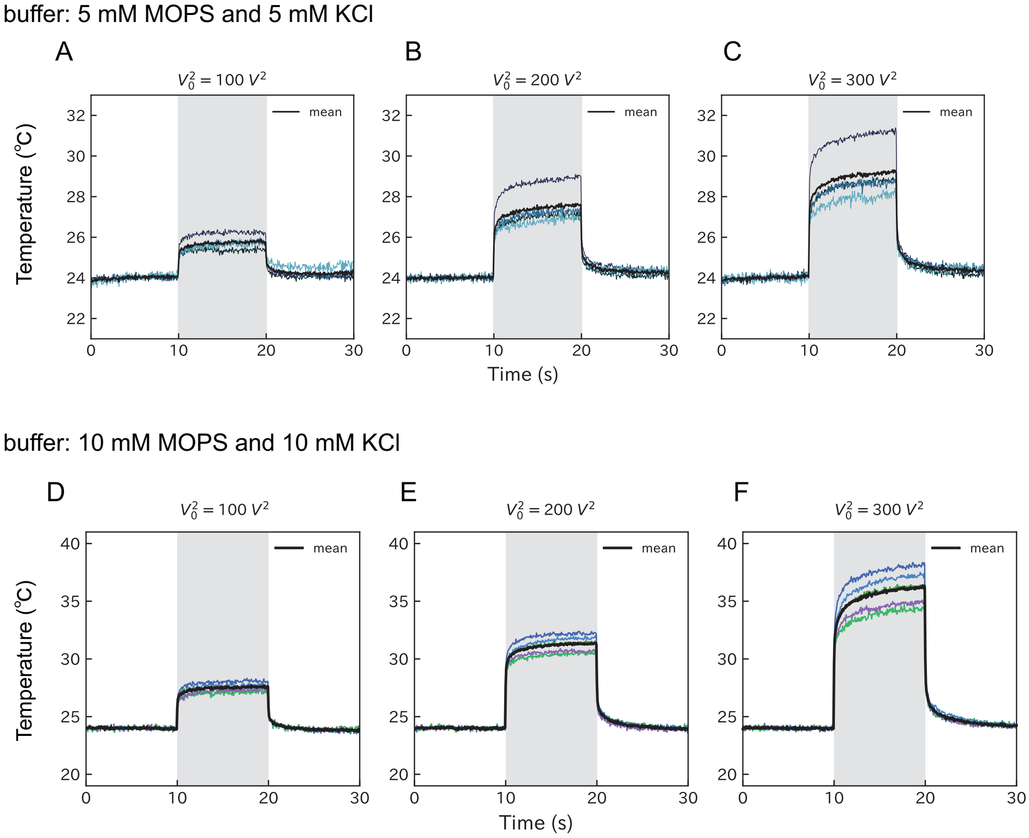


**Figure S2.** Temperature rise by the induction of the electrorotation in the observation buffer containing 5 mM MOPS and 5 mM KCl (**A**–**C**) or 10 mM MOPS and 10 mM KCl (**D**–**F**). The electrorotation with the indicated magnitude was applied from 10 s to 20 s. Black thick line indicates the mean.

#### S3. Remodeling under degraded condition

Figure S3 shows the remodeling traces and dwell time of the cells without a shearing treatment, blocking agent with reduced concentration (50 mg/ml Perfect Block instead of 250 mg/ml), and observation buffer containing high ion strength (10 mM MOPS and 10 mM KCl).

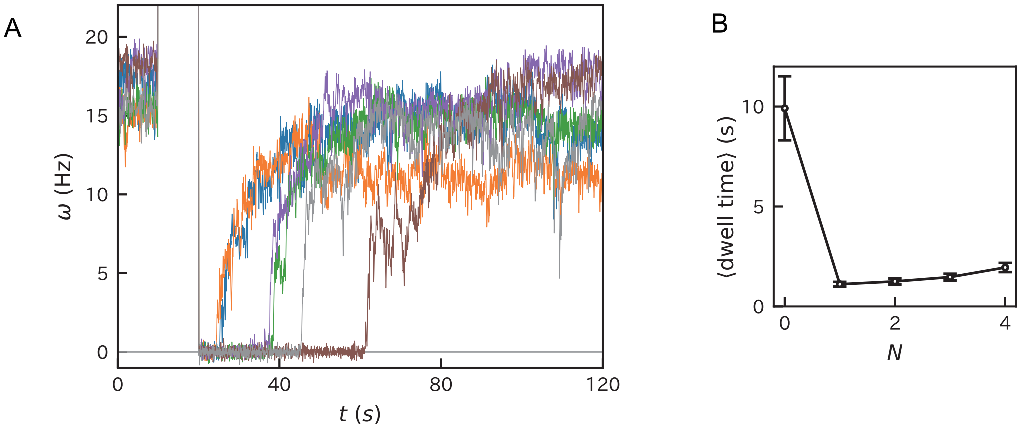


**Figure S3.** Typical remodeling traces (**A**) and dwell time (**B**) under degraded condition. The number of traces observed was 38.

#### S4. Remodeling traces under forced rotation

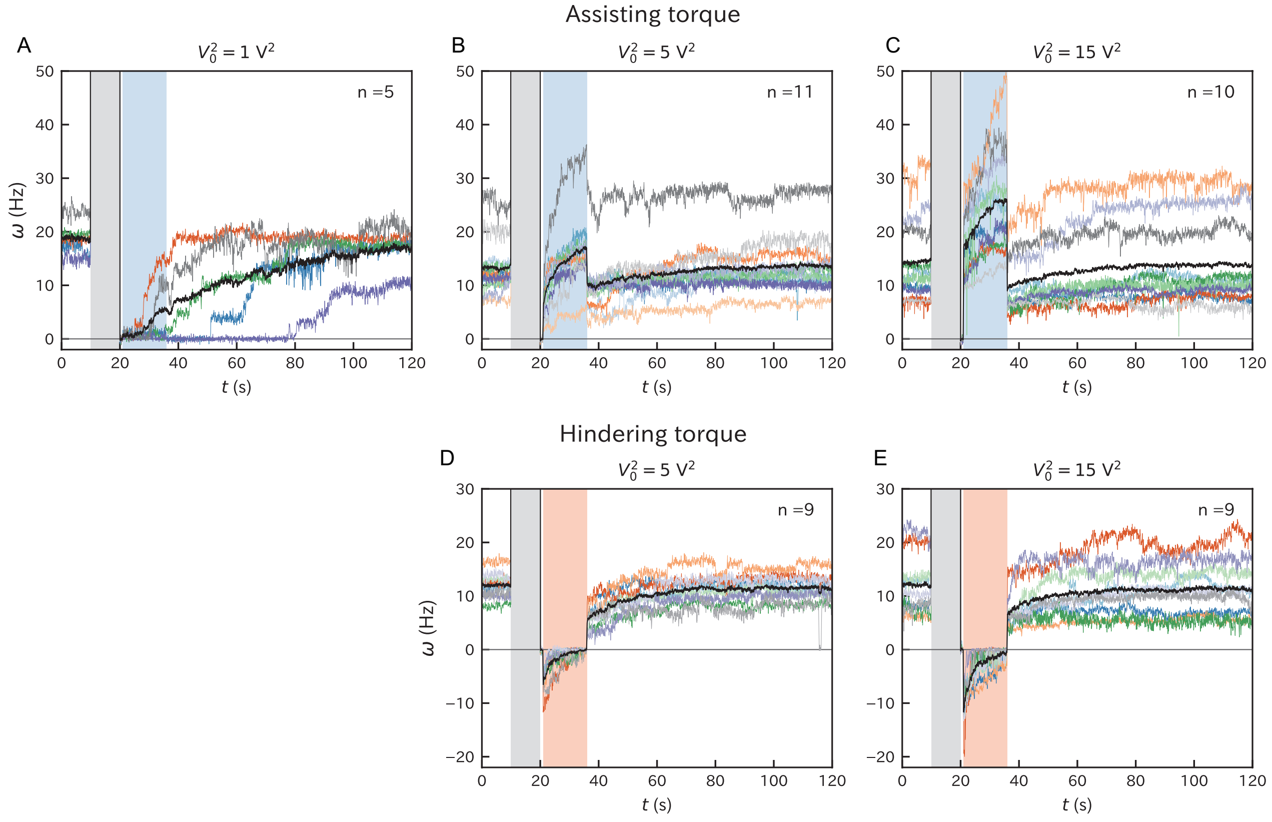


**Figure S4.** Remodeling traces under forced rotation with different torque magnitude. Black curves correspond to the ensemble averages.

#### S5. Quantitative evaluation of rate constant

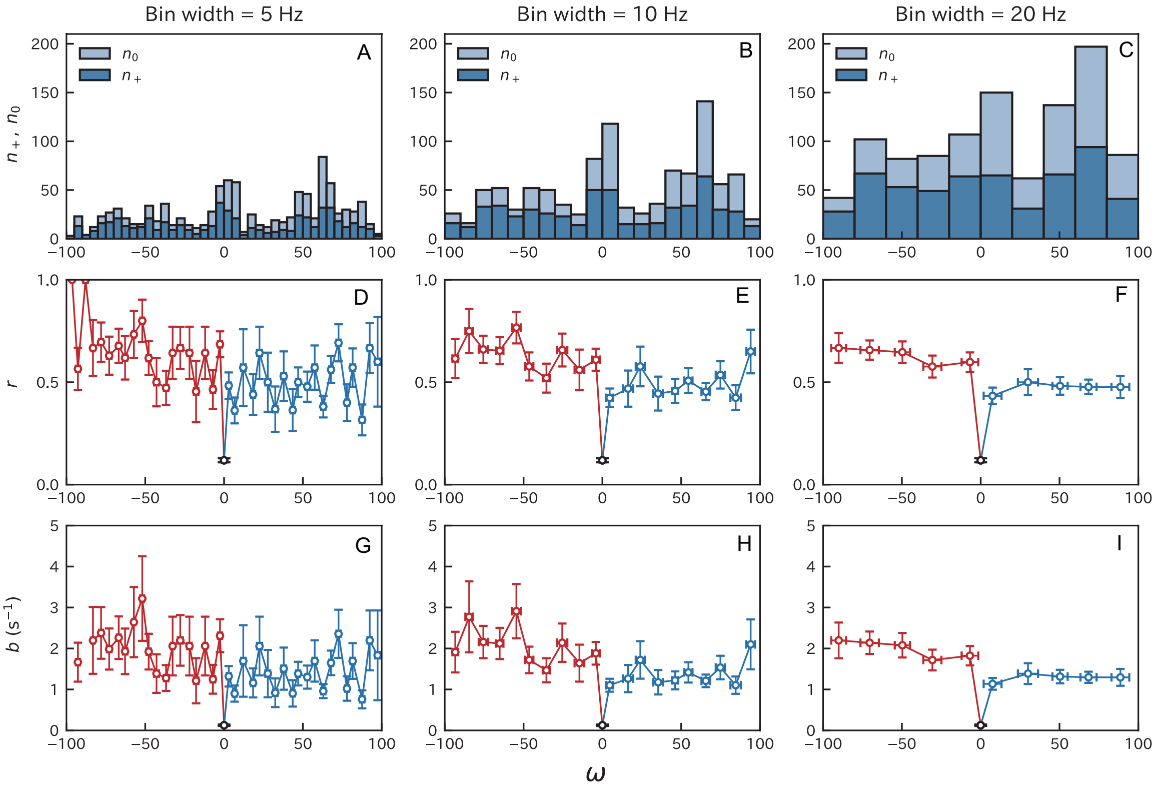


**Figure S5.** Summary of the binding parameters with different bin widths. **A–C**, $n_+$ (blue) and $n_0$ (light blue). **D–F**, $r = n_+/(n_+ + n_0)$. Red and blue correspond to the CW and CCW rotations, respectively. **G–I**, the effective binding rate given by $b = -\tau^{-1} \ln(1-r)$. The error bars correspond to the standard errors.
